# Characterization of cytokine treatment on human pancreatic islets by top-down proteomics

**DOI:** 10.1101/2025.05.12.653562

**Authors:** Ashley N Ives, Tyler Sagendorf, Lorenz Nierves, Tai-Tu Lin, Ercument Dirice, Rohit N Kulkarni, Ljiljana Paša-Tolić, Wei-Jun Qian, James M Fulcher

## Abstract

Type 1 diabetes (T1D) results from autoimmune-mediated destruction of insulin-producing β cells in the pancreatic islet. This process is modulated by pro-inflammatory cytokine signaling, which have been previously shown to alter protein expression in *ex vivo* islets. Herein, we applied top-down proteomics to globally evaluate proteoforms from human islets treated with proinflammatory cytokines (interferon-γ and interleukin-1β). We measured 1636 unique proteoforms across 6 donors and two time points (control and 24-hours post-treatment) and observed consistent changes in abundance across the glicentin-related pancreatic polypeptide (GRPP) and major proglucagon fragment regions of glucagon, as well as the LF-19/catestatin and vasostatin-1/2 region of chromogranin-A. We also observe several proteoforms that increase after cytokine-treatment or are exclusively observed after cytokine-treatment including forms of beta-2 Microglobulin (B2M), high-mobility group N2 protein (HMGN2), and chemokine (C-X-C motif) ligands (CXCL). Together, our quantitative results provide a baseline proteoform profile for human islets and identify several proteoforms that may serve as interesting candidate markers for T1D progression or therapeutic intervention.

**Statement of significance of the study:** This work applies a top-down proteomics workflow for the characterization and label-free quantification of proteoforms from human islets in the context of inflammation. The workflow is optimized for challenges unique to the islet proteome including high disulfide-linkage content and frequent truncation events, resulting in many proteoforms <5 kDa. There are limited examples of top-down proteomics characterization of human islets, thus this study provides a baseline characterization of the proteoforms of major hormones including insulin, glucagon, chromogranin-A and -B, and somatostatin. The quantitative results of proteoform abundances before and after cytokine treatment, which mimics the proinflammatory environment during T1D progression, enables the discovery of interesting proteoform candidates as potential markers of T1D progression.

## 1 Introduction

Type 1 diabetes (T1D) is a devastating disease affecting ∼8.4 million people globally as of 2021, and this burden is expected to increase rapidly.^[1]^ T1D is caused by autoimmune-mediated destruction of insulin-producing β cells in the pancreatic islets.^[2]^ A hallmark of T1D onset is the inflammation of islets (i.e., insulitis) where pro-inflammatory cytokines, including interferon (IFN)-γ and interleukin (IL)-1β, modulate β cell loss.^[3,4]^ A prior bottom-up proteomics study has characterized proteomic changes in islets following IL-1β and IFN-γ treatment as a model for islet inflammation and identified several protecting factors that mediate β cell death.^[5]^ Additionally, several converging studies have demonstrated how prohormone processing is altered in both early and advanced stages of T1D.^[6]^ Top-down proteomics (TDP),^[7]^ which analyzes intact proteins (i.e., without enzymatic digestion), can provide unique insights into prohormone processing products as it captures all mutations, alternative splicing events, proteolytic cleavages, and other post-translational modifications (PTMs). Collectively, these broad classes of protein modifications contribute to the exact molecular form of a protein, or “proteoform”.^[8]^ To date, there have been several top-down imaging mass spectrometry studies of human pancreatic tissue and traditional TDP liquid chromatography-mass spectrometry (LC-MS/MS) studies of mouse islets;^[9–13]^ however, the proteoform composition of human islets is relatively unknown as is the contribution of human islet proteoforms to T1D. We therefore sought to apply TDP to human pancreatic islets in the context of proinflammatory conditions similar to those encountered in T1D.

Herein, we applied an ion-mobility enabled LC-MS/MS TDP analysis on islets from 6 human donors, using both control (i.e., non-treated) and islets exposed to interleukin-1β (IL-1β) and interferon-γ (IFN-γ) for 24 hours as a model of T1D-onset. To increase proteoform coverage, we applied a recently developed high-field asymmetric waveform ion mobility spectrometry (FAIMS) approach that fractionates proteoforms in the gas phase during LC-MS/MS analysis.^[14,15]^ We identified 1636 distinct proteoforms, with 904 proteoforms being quantifiable between controls and cytokine-treatment groups. We measured consistent changes in the abundance of glicentin-related pancreatic polypeptide (GRPP) and major proglucagon fragment regions of glucagon, as well as the LF-19/catestatin and vasostatin-1/2 region of chromogranin-A following cytokine-treatment. We also observed several individual proteoforms that increase after cytokine-treatment or are exclusively observed after cytokine-treatment including forms of beta-2 Microglobulin (B2M), high-mobility group N2 protein (HMGN2), glucagon (GCG), and chemokine (C-X-C motif) ligands (CXCL). Overall, our quantitative results provide a baseline proteoform profile for human islets and identify several proteoforms that may be useful in identifying the onset of T1D or candidate targets for therapeutic intervention.

## 2 Materials and Methods

### 2.1 Experimental Design and Statistical Rationale

Human islets from six non-diabetic cadaveric donors were obtained from the Integrative Islet Distribution Program (IIDP). ∼150 islets per condition were cultured in 2 mL Standard Islet Medium (Prodo) supplemented with human AB serum (Prodo), Ciprofloxacin (Fisher), and glutamine and glutathione (Prodo) at 37 °C under 100% humidity and 5% CO_2_. Islet cultures were allowed to acclimate overnight and then were either treated with cytokines IL-1β and IFN-γ by adding fresh medium containing 50 U/mL and 1000 U/mL of IL-1β and IFN-γ, respectively, or left untreated by adding fresh medium without cytokines for 24 hr. Because the tissues came from cadaveric donors, the study was not considered human subjects research, and no consent was required. The characteristics of the tissue donors are listed in Supplementary Table 1. Prior to data collection, the 12 samples (6 donors, two time points per donor) were randomized. Each sample was analyzed once. For data analysis, samples from the same donor were paired when implementing the limma package^[16]^.

### 2.2 LC-MS/MS Sample Preparation

Islets were resuspended in 466 µL of homogenization buffer (8 M urea, 100 mM ammonium bicarbonate, 5 mM EDTA) and briefly vortexed prior to heating at 37°C for 30 minutes. Reduction was accomplished through addition of 14 µL of 0.5 M dithiothreitol (DTT) with incubation at 20°C for 2 hours within a ThermoMixer (ThermoFisher) set at 1200 RPM. This was followed by alkylation using 40 µL of 0.25 M iodoacetamide (IAA) and incubation at 20°C for 1 hour within a ThermoMixer (ThermoFisher) set at 1200 RPM. The alkylation reaction was quenched with the additin of 50 uL of 0.5 M DTT. Samples were then clarified via 15 minutes of centrifugation at 18,000 RCF at 16 °C. The resulting supernatant was added to a 3 kDA MWCO Amicon® Ultra 0.5 mL centrifugal filter (MilliporeSigma) and centrifuged at 14,000 RCF for 60 minutes at 16°C. ∼50 µL of retentate was then diluted with 0.5 mL wash buffer (8 M urea, 10 mM ammonium bicarbonate, 2 mM EDTA) followed by centrifugation at 14,000 RCF for 60 minutes at 10 °C. This step was performed once more to ensure >100-fold dilution of reduction and alkylation reagents. The concentration of retentates was then determined in duplicate via bicinchoninic acid assay using bovine serum albumin calibration standards prepared in wash buffer (8 M urea, 10 mM ammonium bicarbonate, 2 mM EDTA). Prior to liquid chromatography– mass spectrometry (LC-MS) analysis, samples were adjusted to equivalent concentrations (0.04 mg/mL) with 4 M urea, 5 mM ABC, 1 mM EDTA, 0.5% FA. To reduce any potential for sample loss due to nonspecific binding to surfaces, samples were added to polypropylene PCR tubes inserted into LC-MS vials.^[17]^

### 2.3 LC-MS/MS

Samples were analyzed using a Waters NanoACQUITY UPLC system with mobile phases consisting of 0.2% FA in H2O (Mobile Phase A) and 0.2% FA in acetonitrile (ACN) (Mobile Phase B). Both trapping column (150 µm i.d., 5-cm length) and analytical column (100 µm i.d., 50-cm length) were slurry-packed with C2 packing material (5 µm and 3 µm for trap/analytical respectively, 300 Å, Separation Methods Technology). Samples were loaded into a 10-µL loop, corresponding to 400 ng of loaded material, and injected into the trapping column with an isocratic flow of 5% B at 5 µL/min over 10 minutes for desalting. Separation was performed by ramping from 5 to 15% MPB over 1 minute, followed by 15 to 90% MPB over 89 minutes. The flow rate was maintained at 300 nL/min.

For MS/MS analysis of proteins, the NanoACQUITY system was coupled to a Thermo Scientific Orbitrap Fusion™ Lumos™ Tribrid™ mass spectrometer equipped with the FAIMS Pro interface. Source parameters included electrospray voltage of 2.2 kV, transfer capillary temperature of 275 °C, and ion funnel RF amplitude of 30%. FAIMS parameters were set as previously described^[11]^. FAIMS was set to standard resolution without supplementary user-controlled carrier gas flow and a dispersion voltage (DV) of −5 kV (equivalent to a dispersion field of −33.3 kV/cm), while the CV switched between 3 voltages (−55, −45, and −35) throughout data collection. The Fusion Lumos was set to “Peptide” acquisition mode, and data were collected as full profile. MS1 and MS2 data were acquired at a resolution of 120k and 60k, 2 microscans across a 500 to 2000 m/z range, and with Automatic Gain Control (AGC) targets of 1E6 and 5E5, respectively. MS1 and MS2 were acquired with a maximum inject time of 250 ms. Data dependent settings included selection of top 6 most intense ions, exclusion of ions lower than charge state 3+, exclusion of undetermined charge states, and dynamic exclusion after 1 observation for 30 seconds. Ions selected for MS2 were isolated over a ±1.5 m/z window and fragmented through collision-induced dissociation (CID) with a normalized collision energy of 35%.

### 2.4 Proteoform Identification and Statistical Analysis

Proteoform identification was performed with TopPIC version 1.7.3.^[18]^ Settings for TopPIC included a precursor window of 3 m/z (to account for isotopic envelope), mass error tolerance of 15 ppm, a proteoform cluster error tolerance of 0.8 Da, a mass shift upper bound of 4000 Da and lower bound of −150 Da, and a maximum number of allowed unknown modifications of 1. MS2 spectra were searched against the Swiss-Prot database for *Homo sapiens* containing 20,371 reviewed entries, a variably spliced (“VarSplic”) database containing 21,980 splice-isoform entries, and a TrEMBL database containing 57,749 entries (UP000005640-accessed September 15th, 2022). All databases were scrambled to generate decoys which were concatenated during the search. Carbamidomethylation at cysteine was listed as a static modification. A list of 14 dynamic modifications (N-terminal methionine excision; N-terminal acetylation, acetylation at lysine; C-terminal amidation; N-terminal carbamylation; carbamylation at lysine; deamidation at glutamine or asparagine; methylation or dimethylation at lysine or arginine; iron adduction at aspartic or glutamic acid; phosphorylation at serine, threonine, or tyrosine; N-terminal pyroglutamate; and dioxidation or oxidation at cysteine, methionine, or tryptophan) were provided during the open modification search to reduce the number of unknown mass shifts. The spectra confidence threshold in the TopPIC searches was set at maximum allowable E-Value 0.05.

Downstream data analysis steps were performed in the R environment for statistical computing and TopPICR package.^[19]^ Associated code and functions are provided via Github (https://github.com/ashleyives/top_down_islets_cytokine). Proteoform spectrum matches were filtered to achieve a 1% false discovery rate (FDR). To be considered for differential expression analysis, proteoforms must be observed in both control and treatment groups for at least two patients. For calculation of log2 fold-change, label-free quantities were normalized using two-way median polishing via the medpolish function in R. Differential expression analysis was then performed using the limma R package.^[16]^ P values were generated using a moderated t-test and adjusted for multiple tests using the Benjamini and Hochberg procedure. Manual MS/MS validation was performed using TDValidator v1.125084.1 (Proteinaceous, Inc.; Evanston, IL). MS/MS data was matched to candidate proteoform sequences using the following parameters: 10 ppm tolerance, 3 ppm sub tolerance, 0.35 ppm cluster tolerance, S/N tolerance ≥3, fitter score of ≥0.5 for terminal fragment ions. Theoretical fragment ions were generated using BRAIN.^[20]^

## 3 Results

### 3.1 Coverage of the Proteome

Islets from 6 human donors were divided into control and treatment groups. The treated islets were exposed to interleukin-1β (IL-1β) and interferon-γ (IFN-γ) for 24 hours (**Figure 1A**). After LC-MS/MS analysis and downstream data processing, 874 ±119 (mean±s.d.) proteoforms and 223±34 genes were detected per dataset (**Figure 1B**). Overall, 1636 distinct proteoforms derived from 295 genes could be detected across all 12 datasets (**Figure 1C**). Of these proteoforms, 762 are observed in at least 50% of samples (**Figure 1D**). The mean and median observed monoisotopic masses are 5.7 and 4.1 kDa (**Figure S1**). The RSD distributions and median RSD for the treatment and control groups are modestly high relative to prior TDP analyses on mouse islets. Given mice are raised under more controlled conditions, the additional variance in human samples may be a result of biological differences related to sex, post-mortem interval, or other factors (**Figure S2**).

**Figure 1.**
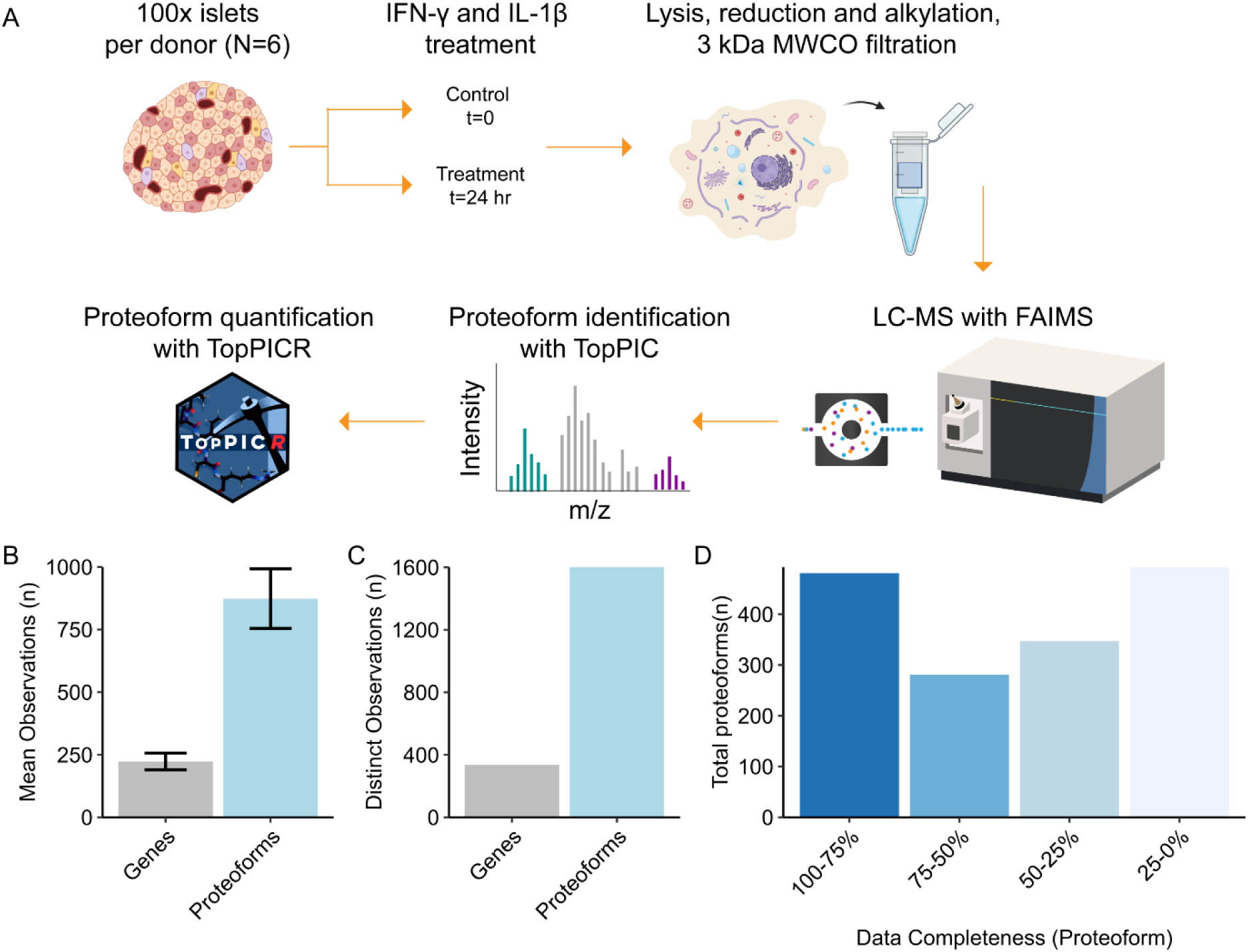
(A) Workflow for processing human islets for top-down proteomic analysis. Created with BioRender.com. (B) Mean number of unique genes (gray) and proteoforms (blue) observed from each human subject (N=6, t=2). Error bars represent +/- sd. (C) Total unique genes (gray) and proteoforms (blue) found in the entire study. (D) Percentile bins containing proteoforms with different degrees of data completeness, e.g. the percentage of acquisitions in which a given proteoform is observed.

### 3.2 Proteoforms of Major Hormones

The most abundant hormones by spectral counting included insulin (INS), glucagon (GCG), chromogranin-A (CHGA), and chromogranin-B (CHGB). Note that we refer to proteoforms in the manuscript by gene and the starting/ending amino acid relative to the full-length protein sequence (i.e. the first amino acid of the signal peptide is residue 1). The most abundant insulin proteoforms included the canonical A- and B-chain (**Figure 2A-B**). We observed various partially processed insulin proteoforms including proinsulin (INS_25-110_), des-64,65 proinsulin (INS_25-87_), des-31,32 proinsulin (INS_57-110_), and proteoforms with non-canonical termini, especially of the C peptide which is detected with both additional N-terminal truncation (INS_58-XX_) as well as extensive C-terminal truncation (INS_57-79/80/81_). We also observed other post-translational modifications including oxidation in both the A- and B-chains. By manual MS/MS validation we localized these oxidations to Cys31, 43, and 108 (**Figure S3**). We also observed pyroglutamylation of the N-terminal glutamate of the C-peptide, along with various water loss, ammonia loss, and iron adduction which are known artifacts of electrospray ionization. We also observed two unknown modifications of insulin including a 128.08 Da mass shift that can be partially localized to the N-terminus of the C peptide and a 13.98 Da mass shift partially localized to the C-terminus of the A-chain. Cross-referencing the UniMod database as well as the expected protein primary sequences in these regions, these additional masses may be the result of N-terminal lysinyl-/glutamylation and the addition of a carbonyl (+O, -2H), respectively. Protein carbonylation is an irreversible oxidation associated with oxidative stress and aging;^[21]^ carbonylation of plasma proteins has been associated with obesity and type 2 diabetes mellitus.^[22]^

**Figure 2.**
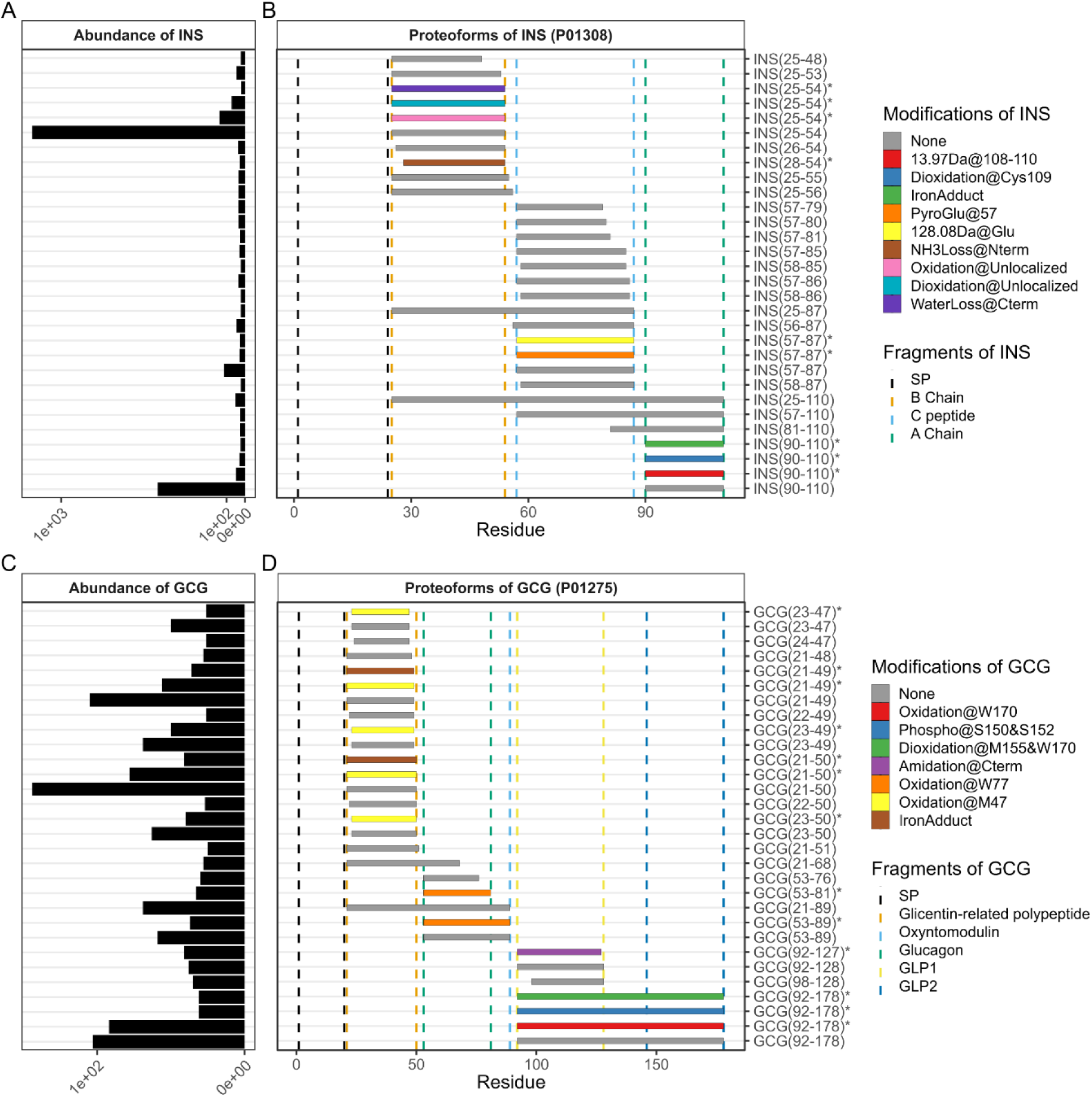
Summary of top 30 most abundant INS and GCG proteoforms. (A) Median spectral count abundance for INS proteoforms. (B) Plot of INS proteoform truncations and modifications (C) Median spectral count abundance for GCG proteoforms. (D) Plot of GCG proteoform truncations and modifications. Right panels map the first and last amino acid of a given proteoform (x-axis), and color fill denotes identified PTMs. Dashed vertical lines annotate the region of a given gene. Proteoforms are sorted top to bottom by ascending C-terminal amino acid ending position, followed by ascending N-terminal amino acid starting position. Y-axis labels denote the first and last amino acid of a given proteoform, and “*” is used to denote modified proteoforms.

The most abundant glucagon proteoforms included the unmodified GRPP (GCG_21-50_), major proglucagon fragment (GCG_92-178_) with and without oxidation, as well as the C-termianlly truncated GRPP (GCG_21-49_) (**Figure 2C-D**). The canonical glucagon fragment (GCG_53-81_) is observed in the top 30 most abundant proteoforms, but as an oxidized form. We observed many truncated proteoforms and oxidation events that cover the glicentin region (GCG_21-89_), which can be further divided into GRPP (GCG_21-50_) and oxyntomodulin (GCG_53-89_); these oxidations are localized to various tryptophan and methionine residues (Met47, Trp77, Met155, and Trp170). We also observed phosphorylation within the proglucagon fragment that is partially localized to either Ser150 and/or Ser152. Finally, we observed three abundant proteoforms of glucagon-like peptide-1 (GLP-1) which include the canonical GLP-1 (GCG_98-128_), a known C-terminally cleaved and amidated form of GLP-1,^[23,24]^ and a known N-terminally cleaved GLP-1 (7-37) (GCG_98--128_)^[23]^.

In addition to INS and GCG, secretory granule-associated chromogranin and somatostatin proteoforms were frequently observed across treated and untreated groups. Observed chromogranin-derived proteoforms generally spanned the entirety of the gene (**Figure S4**), with the most abundant proteoforms including vasostatin-1 (CHGA_19-94_), EA-92 (CHGA_134-225_), amidated GR-44 (CHGA_413-456_), CHGB_21-82_, CHGB_21-83_, and CHGB_21-86_. The most abundant somatostatin proteoforms included somatostatin-14 (SST_103-116_) with and without oxidation, somatostatin-28 (SST_89-116_), prosomatostatin (SST_25-116_), and the N-terminal cleavage product following somatostatin-14 formation (SST_25-100_) (**Figure S5**).

### 3.3 Differential Abundance of Proteoforms Following Cytokine Treatment

We next looked into differential proteoform abundance between the control and cytokine-treated groups. As a criterion for quantification, proteoforms were only considered if they were observed in both control and treatment groups for at least two patients. Therefore, of the 1623 identified proteforms, 904 were considered quantifiable. Applying an unadjusted P value cutoff of < .05 and log2 fold-change cutoff of ±1, 46 proteoforms were found to be decreased in abundance, including several proteoforms of insulin, glucagon, and H1-4 (**Figure 3A-3B, Supplementary File 1**). 39 proteoforms increase in abundance, with the most significant hits including beta-2 microglobulin (B2M), a C-terminally amidated GCG_92-127_ containing a 13.98 Da modification that we tentatively assigned as a carbonyl modification (**Figure S6**), and a truncated form of high mobility group nucleosome-binding domain-containing protein 2 (HMGN2_28-89_). We observed many truncated forms of HMGN1 and HMGN2 (**Figure S7**), however only HMGN2_28-89_ met the cutoff thresholds.

**Figure 3.**
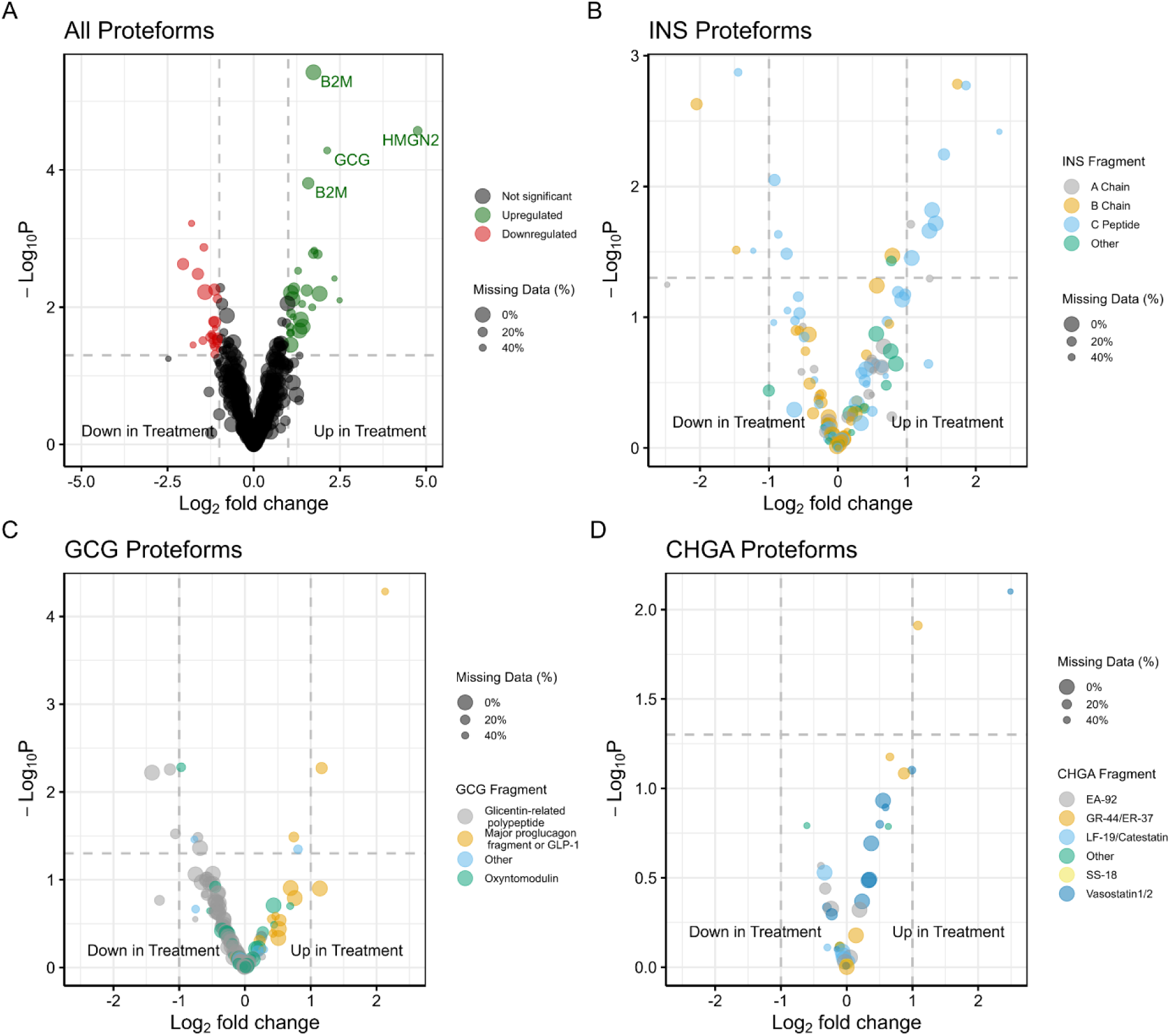
Volcano plots comparing proteoform fold changes pre- and post-cytokine treatment. (A) Volcano plot showing all proteoforms quantified. Proteoforms that are also significant after P value adjustment are annotated with gene names. Color fill denotes if a proteoform has a P value < 0.05 and log2 fold-change cutoff of >1 (green) or <-1 (red), else proteoforms are colored black. (B) Volcano plot showing quantified INS proteoforms. Color fill denotes which region of INS a proteoform is derived from. “Other” denotes proteoforms that span multiple regions. (C) Volcano plot showing quantified GCG proteoforms. Color fill denotes which region of GCG a proteoform is derived from. (D) Volcano plot showing quantified CHGA proteoforms. Color fill denotes which region of CHGA a proteoform is derived from. Horizontal dotted line indicates P value cutoff (.05), and vertical dotted lines indicate log2 fold-change cutoff of 1 and −1. Point size is scaled to the number of missing values present (i.e. larger point size indicates fewer missing values for a given proteoform).

There are also consistent regional trends of altered abundance in GCG (**Figure 3C**) and CHGA (**Figure 3D**). For GCG, 14 out of 16 proteoforms that occur within the major proglucagon fragment (i.e. between amino acids 92 to 178) increased in abundance upon cytokine treatment. Additionally, 71 out of 78 proteoforms that occur within the GRPP fragment (i.e. amino acids 18 to 52) decreased in abundance upon cytokine treatment. For CHGA, 5 out of 6 proteoforms that occur within the LF-19/catestatin fragment (i.e. between amino acids 358 to 390) decreased in abundance upon cytokine treatment. All of these proteoforms begin at amino acid 358 (the LF-19 starting position) and end at residues 376, 378, 386, 389, or 390. Additionally, 9 of 13 proteoforms that occur with the vasostatin-1/2 region (i.e. between amino acids 19 to 131) increase upon cytokine treatment.

Outside of differential abundance, we also wondered if any proteoforms were unique to a given group. As proteoforms in a single group cannot be quantified through differential abundance, we estimated the probability of a proteoform being enriched for either group using a hypergeometric probability distribution. Through this analysis, we observed several proteoforms all belonging to the chemokine (C-X-C motif) ligand (CXCL) family that are unique to the cytokine treatment condition (**Figure 4**). This includes various truncated forms of CXCL1, CXCL9, CXCL10, and CXCL11. Several of these proteoforms are observed across all patients in the treatment condition, including CXCL1_35-107_, CXCL10_22-98,_ CXCL10_25-94,_ and CXCL10_22-94_. We did not observe statistically significant proteoforms unique to the control group.

**Figure 4.**
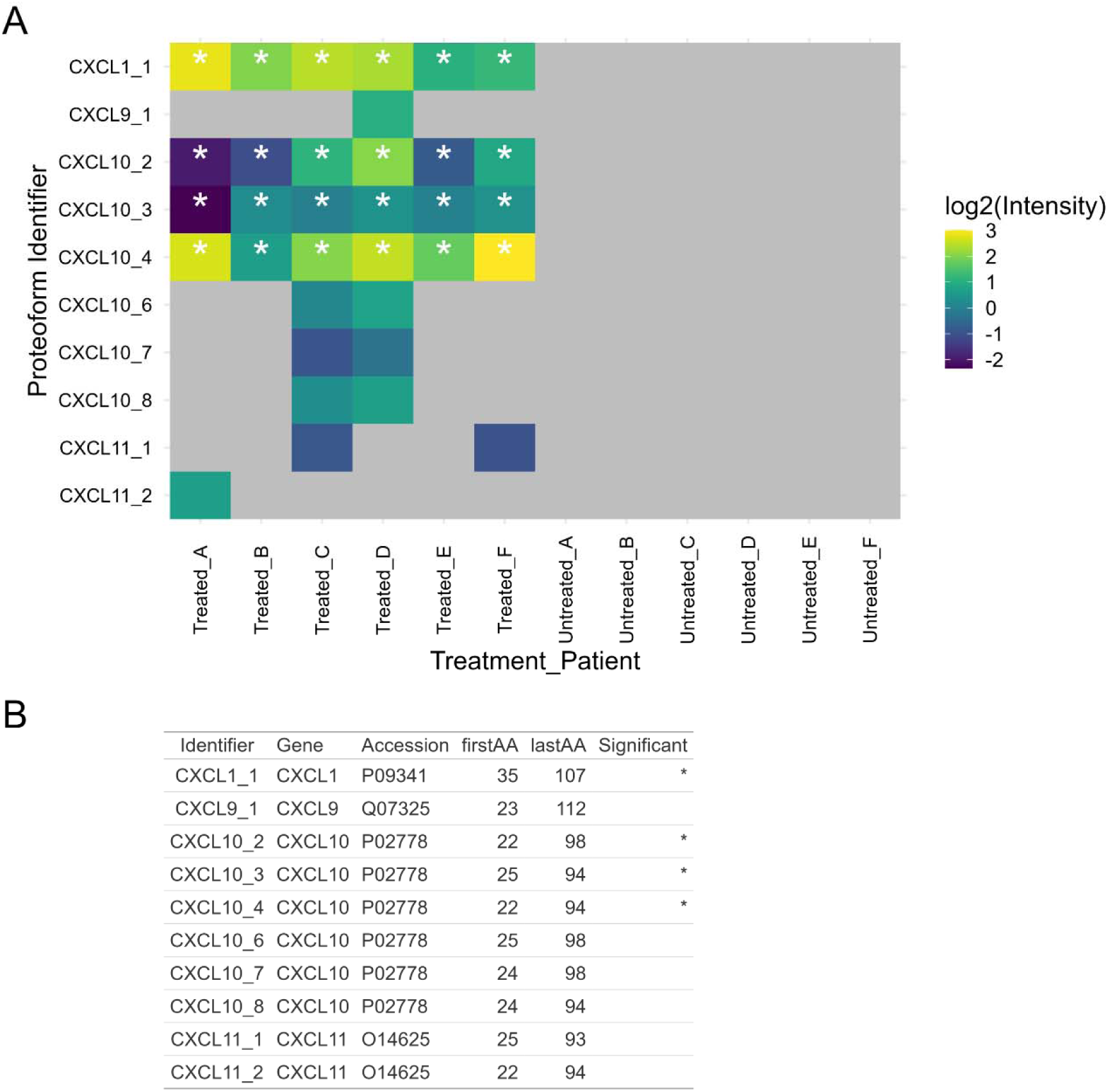
Proteoforms unique to cytokine treatment condition. (A) Heatmap of proteoforms unique to the cytokine treatment condition. Data is plotted as arbitrary proteoform identifiers (Gene_##) versus treatment_patient identifier. Fill denotes the median normalized log2(Intensity) as determined by label-free quantification. NA values are shown in gray. Asterisks denote proteoforms that are significant (probability < 0.01) using a hypergeometric probability distribution. (B) Summarizes all proteoforms plotted in panel (A) including the first and last amino acid (firstAA, lastAA) based on the listed UniProt Accession.

## 4 Discussion

Despite the recognized role of altered prohormone processing in T1D,^[6]^ there is limited molecular-level proteoform data for hormones from healthy or metabolically stressed human-donors. Herein, we have applied top-down proteomics to human islets stressed with proinflammatory cytokines. Overall, we observed many oxidation events at cysteine, tryptophan, methionine, and possibly other carbonylated residues which are known byproducts of reactive oxygen species (ROS).^[25]^ We also observed shifts in proteoform abundance for the GLP and GRPP regions of glucagon as well as the LF-19/catestatin and vasostatin regions of chromogranin-A upon cytokine treatment. Our observation that GLP-1 and major proglucagon fragment (GCG_92-178_) proteoforms increase upon cytokine treatment agrees with prior reports of pre-diabetic individuals have elevated levels of secreted of GLP-1 peptide,^[26]^ possibly due to adaptive alterations in prohormone convertase (PC) enzyme activities. Less is known about glicentin and GRPP dervied proteoforms, as these hormones do not have a known receptor and the GRPP region is poorly conserved in mammals.^[27,28]^ However, monitoring GRPP proteoforms may be of future interest given that GRPP inhibits insulin secretion in rodent models.^[29]^ For CHGA derived proteoforms, it is generally understood that circulating levels of CgA (CHGA_19-457_) and pancreastin (CHGA_272-319_) are elevated in T1D subjects^[30]^ and that several CHGA derived peptides (including vasostatin-1) are autoantigens for β-cell-destructive diabetogenic T-cells.^[31–33]^ The exact impact of altered LF-19/catestatin abundance is currently unclear, however catestatin has also been shown to impact hepatic glucose production and insulin sensitivity in rodent models.^[34]^

A unique advantage of top-down proteomics is that ability to characterize truncated forms of a given gene product.^[35]^ We observed many non-canonical truncated forms of the major hormones including insulin, glucagon, chromogranin-A, chromogranin-B, and somatostatin, demonstrating the complexity of hormone processing products that can only be visualized by intact protein analysis. Additionally, many chemokines are proteolytically cleaved as a means to regulate their chemotactic function; these cleavages can both activate or inhibit chemotactic activity or chemokine receptor selectivity.^[36]^ We consistently observed CXCL1 and CXCL10 across all donors, which are known to be released following exposure to IL-1β^[37]^ and IFN-γ.^[38–40]^ We observe the full length CXCL10 with the signal peptide excised (CXCL10_22-98_), as well as C-terminally truncated CXCL10_22-94_ (alternatively known as CXCL10[1-73]) which is a less potent ligand using CXCR3A-mediated signaling assays.^[41]^ CXCL1 and CXCL10 have been found to be elevated in the serum levels of T1D subjects,^[42,43]^ underscoring the importance of characterizing these proteoforms. Other no0 truncated proteoforms included an unpregulated form of HMGN2 which is cleaved in nucleosome binding domain as well at the final C-terminal Lys residue (HMGN2_28-89_).^[44,45]^ Interestingly, the nucleosome binding domain of HMGN2 is highly conserved and consists of a 30 amino acid long region that is highly basic.^[46]^ Considering the high carboxypeptidase activity in islets targeting basic residues such as Lys or Arg, it is possible that the basic residues in HMGN2 are a target for carboxypeptidases. We detected other unique truncation states of HMGN2 and HMGN1, however it is unclear from the data coverage if these forms are significantly impacted by cytokine treatment. HMGN3 has previously been linked to regulation of insulin secretion and a diabetic phenotype in mouse models;^[47]^ it is unclear of HMGN1/2 simiarly regulate hormone secretion but the HMGN family and highly conserved and share a large sequence identity.^[48,49]^

Label-free quantification for top-down proteomics is an active area of research, with many data analysis and technical optimizations being developed for improved performance.^[50–52]^ Here, we have applied a top-down proteomics workflow for the characterization and label-free quantification of proteoforms from human islets. This study provides a baseline characterization of the major hormones including insulin, glucagon, chromogranin-A and -B, and somatostatin. Additionally, the proteoforms responsive to cytokine treatment may prove useful for future studies of T1D-specific biomarkers and therapeutic targets. Future application of this pipeline to larger cohorts and cohorts with more diverse metabolic states will offer valuable insights into the relevance and practical application of pancreatic hormone proteoforms in the context of T1D.

## 5 Associated Data

All scripts, functions, and source data are available at the Github repository (https://github.com/ashleyives/top_down_islets_cytokine). The mass spectrometry raw data have been deposited to the ProteomeXchange Consortium via the MassIVE partner repository with dataset accession MSV000097810.

## Acknowledgements

This work is supported by National Institutes of Health grants U01DK137113, R01DK135081, and R01DK122160. We thank Dr. Matthew Monroe for his assistance in depositing the raw proteomic data onto MassIVE. A portion of the research was performed using EMSL (grid.436923.9), a DOE Office of Science User Facility sponsored by the Biological and Environmental Research program.

## Abbreviations

ABC: ammonium bicarbonate
AGC: Automatic Gain Cntrol
DDM: n-Dodecyl-β-D665 Maltoside
DDA: data-dependent acquisition
DTT: Dithiothreitol
EDTA: Ethylenediaminetetraacetic acid
FA: Formic Acid
FAIMS: high Field Asymmetric waveform Ion Mobility Spectrometry
FDR: false discovery rate
HCD: higher-energy collisional dissociation
IAA: Iodoacetamide
IFN-γ: interferon-γ
IL-1β: interleukin-1β
LC-MS/MS: liquid chromatogprahy-tandem mass spectrometry
LFQ: label-free quantification
MS: mass spectromery
T1D: type1 diabetes

