## Supplementary Information for "Characterization of cytokine treatment on human pancreatic islets by top-down proteomics"

**Table S1. Patient meta data.**
Age (years), sex, ethnicity, BMI, and HgbA1c (%) for donors.

| Unique Identifier | Age (years) | Sex | Ethnicity | BMI | HgbA1c (%) |
|---|---|---|---|---|---|
| ADC5225A | 27 | M | Hispanic | 30.7 | 5.2 |
| ADC4231 | 29 | M | White | 32.5 | 5.5 |
| ADIL276 | 51 | F | Hispanic | 30.1 | NA |
| ADIR339 | 44 | F | White | 34.5 | NA |
| ADH3495 | 53 | F | Hispanic | 33.3 | 6 |
| ADH1303 | 50 | F | White | 20.3 | NA |

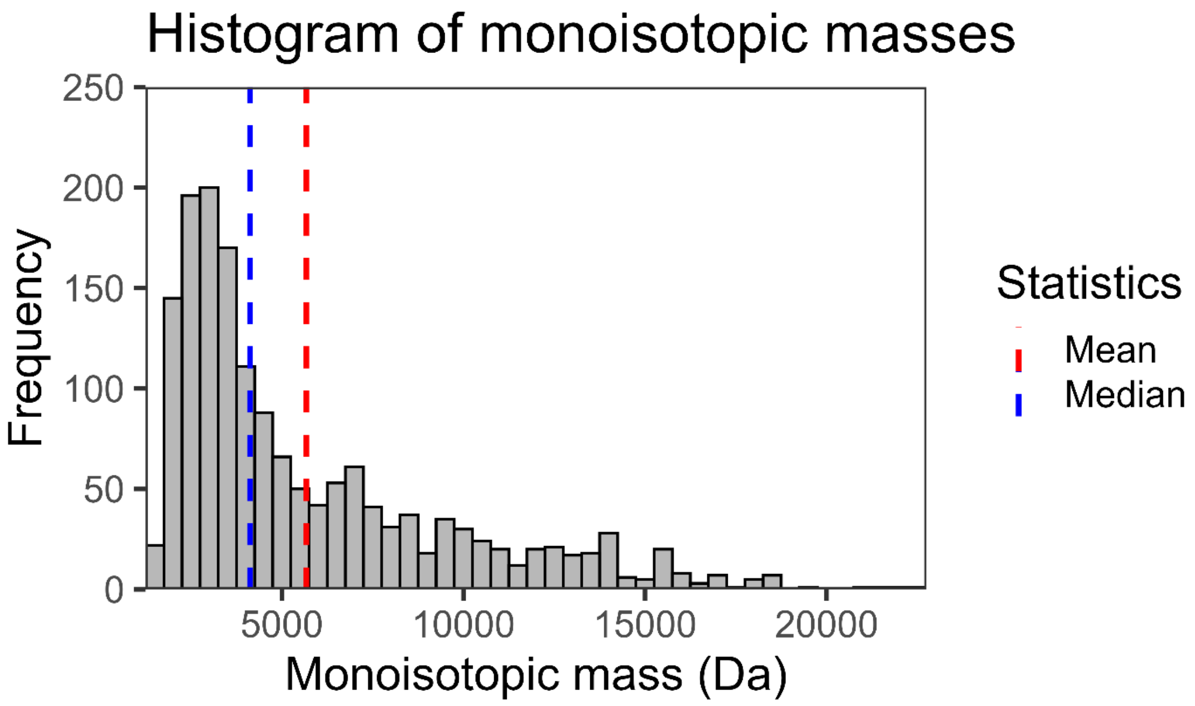

**Figure S1. Histogram of monoisotopic masses for observed proteoforms.**
The mean (red) and median (blue) are annotated as dashed lines.

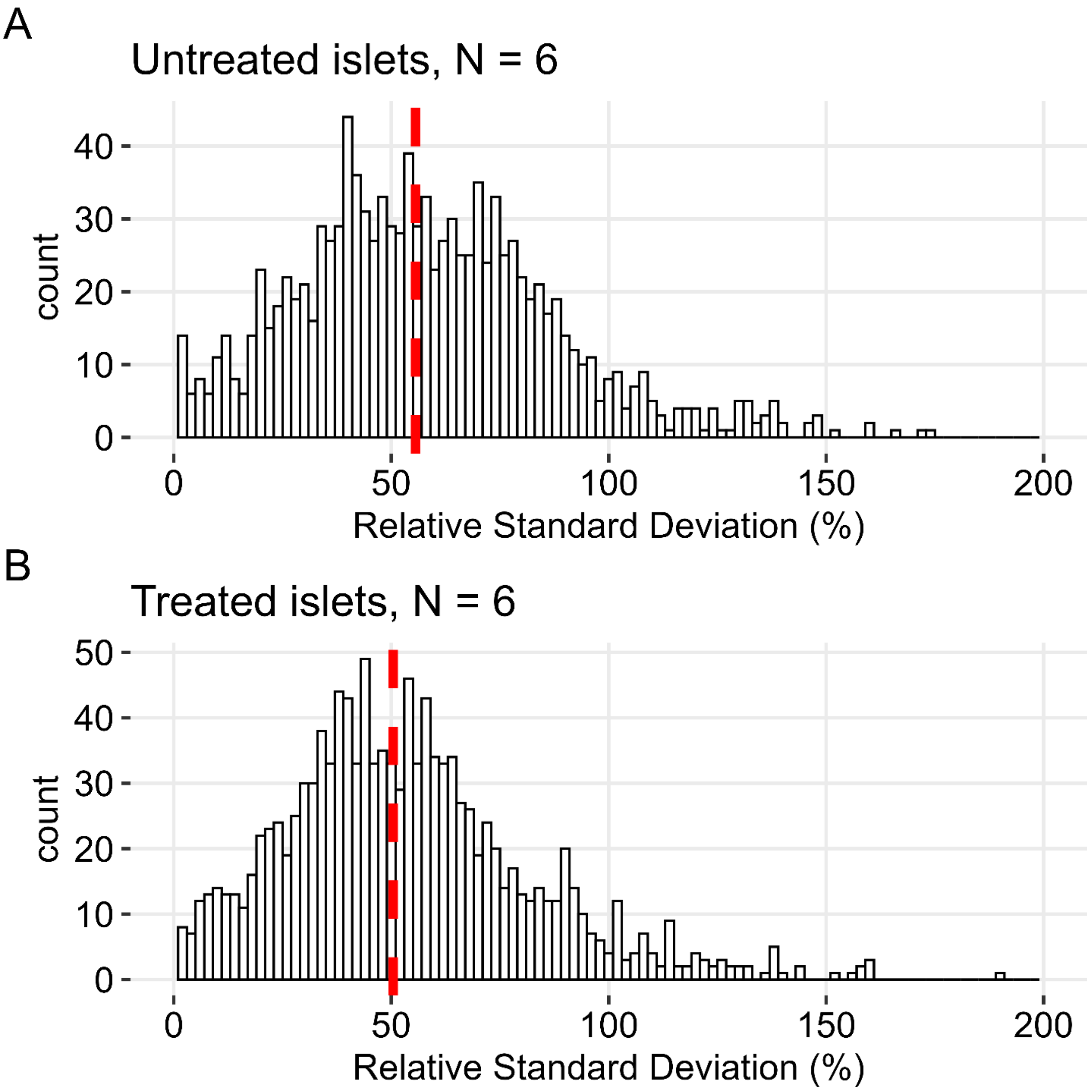

**Figure S2. Histogram of relative standard deviations (RSD) based on label-free, intensity-based quantification of observed proteoforms.**

RSDs are shown for (A) untreated, control islets and (B) islets after cytokine treatment following median normalization. The median (red) is annotated as a dashed line.

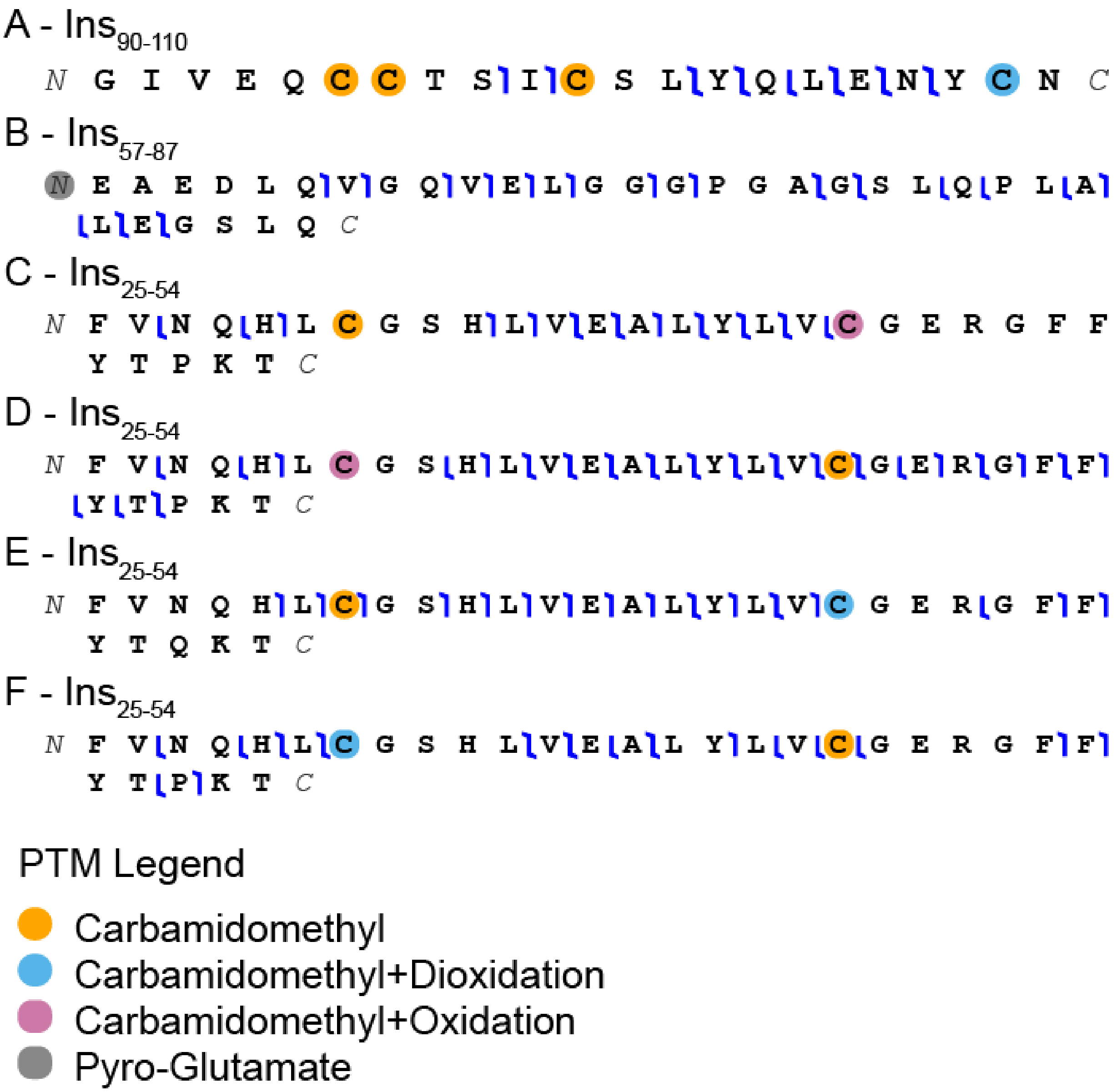

**Figure S3. Fragmentation (MS/MS) maps of select INS proteoforms.**

Maps denote matching fragment ions for searched candidate proteoforms including (A) dioxidation of Cys108, (B) pyro-glutamylation of the C peptide, oxidation within the B chain localized to either (C) Cys43 or (D) Cys31, and dioxidation within the B chain localized to either (E) Cys43 or (F) Cys31. Note that panels (C) and (D) are derived from the same proteoform spectral match scan. (E) and (F) are also derived from the same proteoform spectral match scan. Colored circles denote searched modifications including carbamidomethyl (gold), carbamidomethyl and dioxidation (blue), carbamidomethyl and oxidation (pink), and pyroglutamate (gray). Panel labels denote the first and last amino acid of a searched proteoform (i.e. $Ins_{firstAA\text{-}lastAA}$). Blue flags denote either *b*- (left-facing) or *y*-type (right-facing) fragment ions. The N- and C-termini are denoted as italized letters.

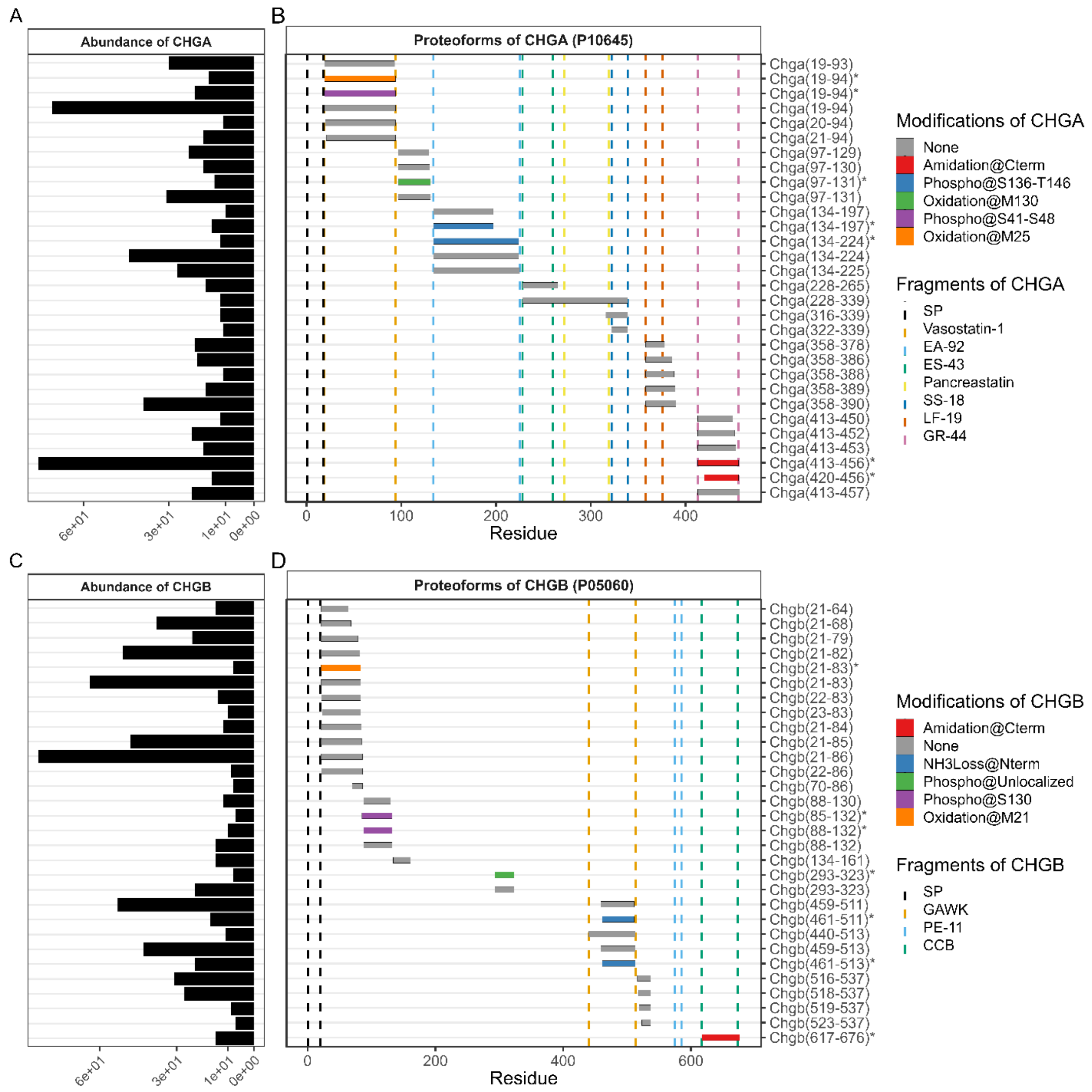

**Figure S4. Summary of top 30 most abundant CHGA and CHGB proteoforms.**
(A) Median spectral count abundance for (B) CHGA proteoforms. (C) Median spectral count abundance for (D) CHGB proteoforms. Rightmost panels map the first and last amino acid of a given proteoform (x-axis), and color fill denotes identified PTMs. Dashed vertical lines annotate the region of a given gene. Proteoforms are sorted top to bottom by ascending C-terminal amino acid ending position, followed by ascending N-terminal amino acid starting position. Y-axis labels denote the first and last amino acid of a given proteoform, and “*” is used to denote modified proteoforms.

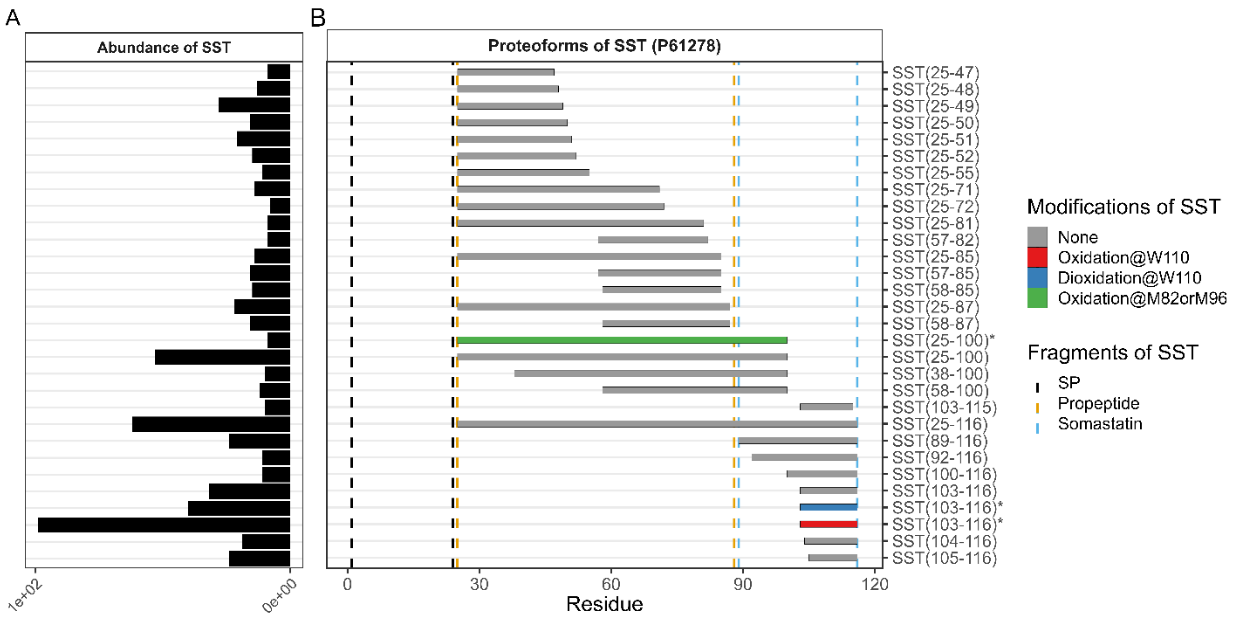

**Figure S5. Summary of top 30 most abundant SST proteoforms.**
(A) Median spectral count abundance for (B) SST proteoforms. Rightmost panels map the first and last amino acid of a given proteoform (x-axis), and color fill denotes identified PTMs. Dashed vertical lines annotate the region of a given gene. Proteoforms are sorted top to bottom by ascending C-terminal amino acid ending position, followed by ascending N-terminal amino acid starting position. Y-axis labels denote the first and last amino acid of a given proteoform, and "*" is used to denote modified proteoforms.

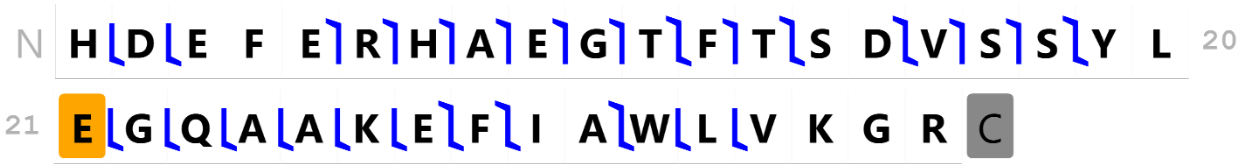

**Figure S6. Fragmentation map for C-terminally amidated, oxidized $GCG_{92\text{-}127}$.**
Assigned fragment ions are shown for the lowest E-value PrSM scan for C-terminally amidated, oxidized $GCG_{92\text{-}127}$. The amidation site is denoted with a gray box. The gold box denotes a carbonyl modification (+O, -2H). Blue flags denote either *b*- (left-facing) or *y*-type (right-facing) fragment ions. The N- and C-termini are denoted as gray letters.

A

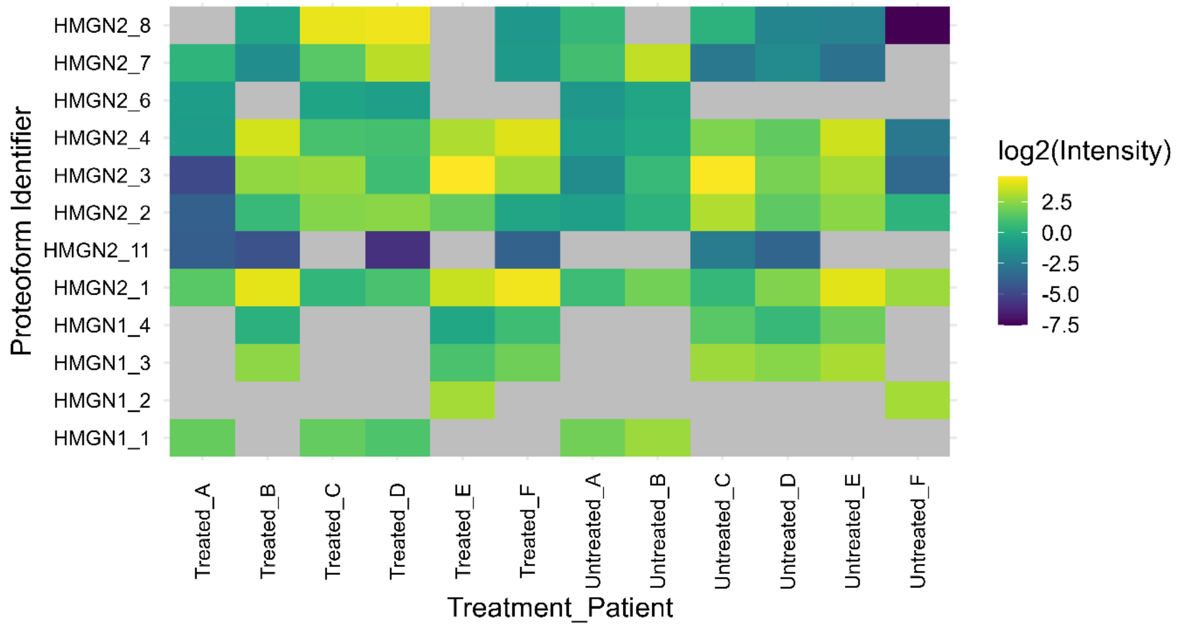

B

| Identifier | Gene | Accession | firstAA | lastAA | Unexpected Modification |
|---|---|---|---|---|---|
| HMGN2_8 | HMGN2 | P05204 | 28 | 89 | FALSE |
| HMGN2_7 | HMGN2 | P05204 | 25 | 89 | FALSE |
| HMGN2_6 | HMGN2 | P05204 | 2 | 78 | FALSE |
| HMGN2_4 | HMGN2 | P05204 | 25 | 90 | FALSE |
| HMGN2_3 | HMGN2 | P05204 | 28 | 90 | FALSE |
| HMGN2_2 | HMGN2 | P05204 | 43 | 90 | TRUE |
| HMGN2_11 | HMGN2 | P05204 | 25 | 67 | FALSE |
| HMGN2_1 | HMGN2 | P05204 | 2 | 90 | FALSE |
| HMGN1_4 | HMGN1 | P05114 | 54 | 100 | FALSE |
| HMGN1_3 | HMGN1 | P05114 | 24 | 100 | FALSE |
| HMGN1_2 | HMGN1 | P05114 | 2 | 100 | FALSE |
| HMGN1_1 | HMGN1 | P05114 | 2 | 100 | TRUE |

**Figure S7. Proteoforms of HMGN1 and HMGN2.**
(A) Heatmap of HMGN1 and HMGN2 proteoforms. Data is plotted as arbitrary proteoform identifiers (Gene_##) versus treatment_patient identifier. Fill denotes the median normalized log2(Intensity) as determined by label-free quantification. NA values are shown in gray. (B) Summarizes all proteoforms plotted in panel (A) including the first and last amino acid (firstAA, lastAA) based on the listed UniProt Accession. Proteoforms with unknown mass shifts are listed in column “Unexpected Modification”.
